# Reduced nephron endowment in the common *Six2-TGC^tg^* mouse line is due to *Six3* misexpression by aberrant enhancer-promoter interactions in the transgene

**DOI:** 10.1101/2023.10.06.561202

**Authors:** Alison J. Perl, Han Liu, Matthew Hass, Nirpesh Adhikari, Praneet Chaturvedi, Yueh-Chiang Hu, Rulang Jiang, Yaping Liu, Raphael Kopan

## Abstract

Lifelong kidney function relies on the complement of nephrons generated during mammalian development from a mesenchymal nephron progenitor cell (NPC) population. Low nephron endowment confers increased susceptibility to chronic kidney disease. We asked whether reduced nephron numbers in the popular *Six2TGC* transgenic mouse line^1^ was due to disruption of a regulatory gene at the integration site or to ectopic expression of a gene(s) contained within the transgene. Targeted locus amplification identified integration of the *Six2TGC* transgene within an intron of *Cntnap5a* on chr1. We generated Hi-C datasets from NPCs isolated from the *Six2TGC^tg/+^* mice, the *Cited1^CreERT2/+^*control mice, and the *Six2TGC^tg/+^*;*Tsc1^+/Flox,2^* mice that exhibited restored nephron number compared with *Six2TGC^tg/+^*mice, and mapped the precise integration of *Six2TGC* and *Cited1^CreERT2^*transgenes to chr1 and chr14, respectively. No changes in topology, accessibility, or expression were observed within the 50-megabase region centered on *Cntnap5a* in *Six2TGC^tg/+^* mice compared with control mice. By contrast, we identified an aberrant regulatory interaction between a *Six2* distal enhancer and the *Six3* promoter contained within the transgene. Increasing the *Six2TGC^tg^* to *Six2* locus ratio or removing one *Six2* allele in *Six2TGC^tg/+^* mice, caused severe renal hypoplasia. Furthermore, CRISPR disruption of *Six3* within the transgene (*Six2TGC*^Δ*Six3CT*^) restored nephron endowment to wildtype levels and abolished the stoichiometric effect. Data from genetic and biochemical studies together suggest that in *Six2TGC,* SIX3 interferes with SIX2 function in NPC renewal through its C-terminal domain.

**Significance:** Using high-resolution chromatin conformation and accessibility datasets we mapped the integration site of two popular transgenes used in studies of nephron progenitor cells and kidney development. Aberrant enhancer-promoter interactions drive ectopic expression of *Six3* in the *Six2TGC^tg^*line which was correlated with disruption of nephrogenesis. Disruption of *Six3* within the transgene restored nephron numbers to control levels; further genetic and biochemical studies suggest that *Six3* interferes with *Six2*-mediated regulation of NPC renewal.

## INTRODUCTION

Three cell populations – the ureteric bud (UB) tip, the nephron progenitor cells (NPCs) and the stroma – engage in reciprocal signaling cascades supporting UB branching morphogenesis (GDNF/RET), NPC self-renewal and niche retention (FGF, WNT^LO^ and BMP), and NPC niche exit and differentiation (WNT^HI^ and BMP)^3^ *in utero*. NPCs generate the nephrons; the balance between their self-renewal and differentiation is a critical determinant of the final nephron number and is dependent on the strength of canonical Wnt signaling^4–7^ modulated in part by age-dependent translatome changes^2^. Nephrogenesis ends synchronously across multiple niches during gestation (human) or a few days postnatally (mice). In contrast with some vertebrates that produce nephrons throughout life^8^, mammals are only capable of limited nephron repair. Thus, the mammalian nephron number (endowment) can only decline with age and is a critical determinant of disease-free survival. In humans, nephron endowment varies by 10-fold^9^, and low nephron number is associated with increased risk of hypertension and chronic kidney disease^10^. Identifying the genetic and molecular mechanisms regulating nephron number establishment could inform therapeutic approaches to support kidney function across the lifespan of vulnerable populations, such as those born prematurely^9^.

Mouse models provide a tractable experimental system for evaluating the effects of genetic background, allelic manipulations, or the environment on nephron number^10^. We complemented published chromatin accessibility (ATAC-seq) datasets^11^ with chromatin conformation capture (Hi-C) in NPCs isolated from mouse strains and mutants with variable nephron numbers, and matching ATAC-seq datasets from NPCs at different developmental timepoints to seek shared factors that might promote high nephron number. Because haploinsufficiency of *Six2* increased nephron numbers^12^, one might predict that reduced *Six2* accessibility would be associated with high nephron number-producing NPCs of the same age. Contrary to this prediction, *Six2* accessibility is enriched in young NPCs^11^. Older NPCs engage in “lineage priming”^11, 13^, opening enhancers that will be used after the mesenchymal to epithelial transition.

The observation that the *Six2TGC^+/tg^* mice^2, 14^ had reduced nephron numbers led us to ask if this effect in the widely used Cre driver mouse line was transgene-dependent or integration site-dependent. We identified the transgene integration site in a *Cntnap5a* intron on chr1 and mapped its architecture. Local topology, accessibility, and transcription were not altered within 50 megabases (MB) centered on the integration site. *Six3* mRNAs is absent from wild type control NPCs but is expressed in *Six2*-expressing NPCs from the *Six2TGC^+/tg^* mice ^2,14^. We determined that expression is due to an anomalous regulatory interaction between a *Six2* distal enhancer and the *Six3* gene within the bacterial artificial chromosome (BAC). We show that CRISPR-mediated disruption of *Six3* exon1/2 splicing and loss of the C-terminal domain in the transgene restored normal nephron numbers in genome-edited *Six2TGC^+/^*^Δ*Six3CT*^ mice. We propose that *Six3*-mediated interference with SIX2 function^4^ underlies this loss of nephrons in this line.

## RESULTS

### *Six2TGC^+/tg^* NPCs exhibit ectopic *Six3* expression due to an anomalous interaction between the *Six3* promoter and a *Six2* distal enhancer within the transgene

The low nephron number *Six2TGC* transgenic mice drew our attention to the observed expression of *Six3* in *Six2*-expressing NPC^2, 14, 15^, (Figure 1A-B). Though we could not detect SIX3 by immunofluorescence with commercially available antibodies, TRAP-seq^2^ revealed that *Six3* mRNA was nearly as abundant as *Six2 mRNA* in the translatome (7.5CPM for *Six3* and 9.4CPM for *Six2* in the polysomal fraction; Table S5 in^2^), consistent with the presence of SIX3 protein^16, 17^. These mice contain a randomly integrated BAC (clone RPCI23-311C1) encompassing an approximately 170kb region of mouse chr17 including the *Six3* gene locus, the *Six2* distal enhancer (*Six2*-DE), and a heavily modified *Six2* locus in which GFP and Cre have replaced part of the *Six2* exon1 coding sequence^18^. We PCR amplified and sequenced the modified *Six2* exon1 region with primers designed based on the BAC and the original report^18^ and found it contains a nonfunctional tTA cassette with 2xpolyA followed by a CMV minimal promoter-driven eGFP-Cre fusion gene in the middle of *Six2* exon 1 (with tTA sequence inserted 133-bp downstream of the *Six2* ATG codon, out of frame; Supplemental Table S1). Given the distance and the presence of two polyA signal sequences between the *Six2* promoter and *tetO-CMVmini-eGFP-Cre*, we surmised that eGFP-Cre is driven by the *tetO-CMVmini* promoter rather than the *Six2* promoter, likely contributing to the mosaicism observed in this line.

**Figure 1.**
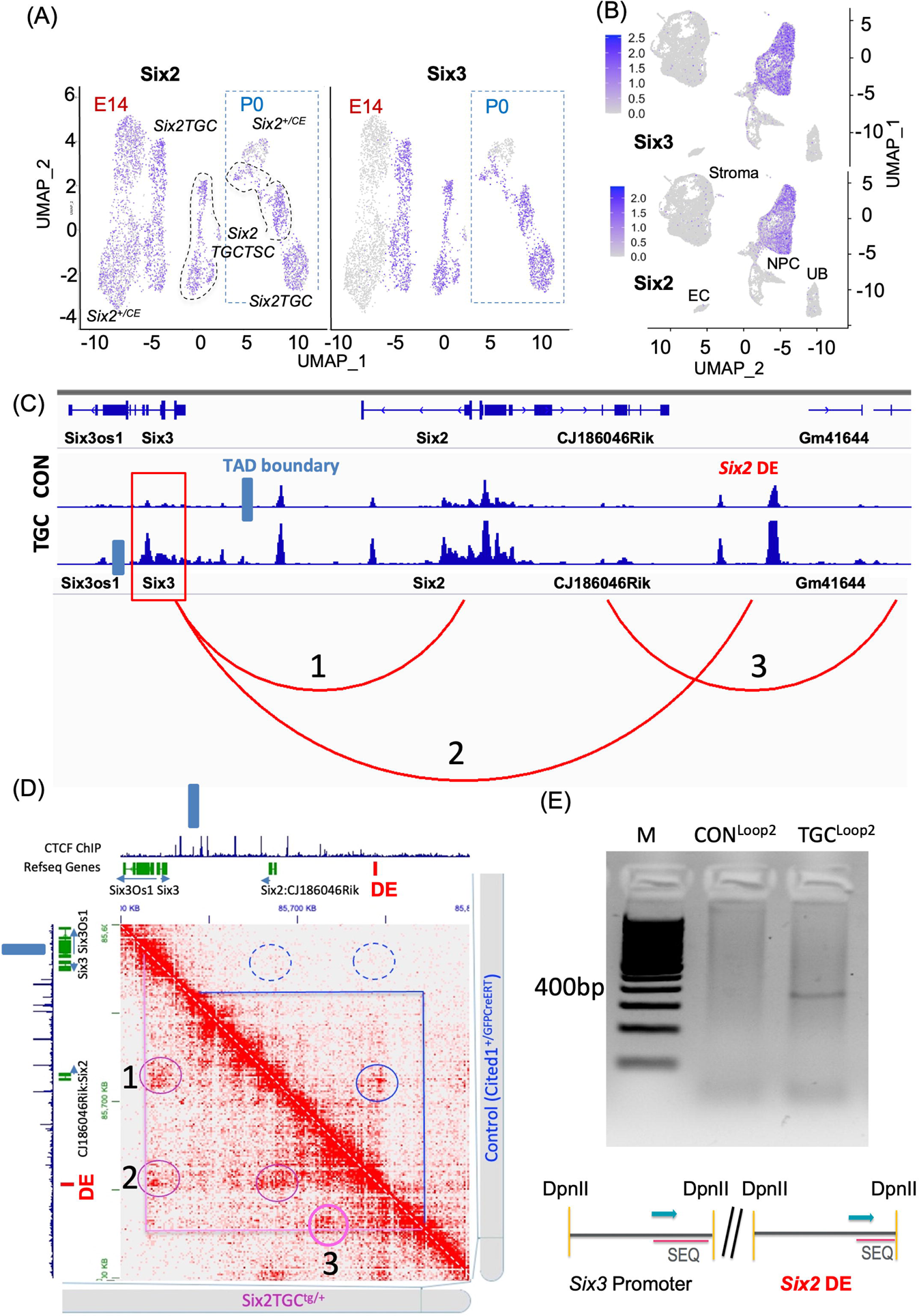
Hi-C and 3C confirm an aberrant regulatory interaction between the *Six3* promoter and *Six2* distal enhancer in *Six2TGC*. (A) UMAP of single-cell RNA-sequencing at E14 and P0 NPCs (reported in^26^ Fig 2L) identifies *Six3* expression in *Six2TGC*-containing NPCs but not in *Six2^+/CE^*NPCs. Expression value show Log2 transform, scaling was performed to normalize the raw counts of genes across all cells. (B) UMAP visualization of *Six3* and *Six2* expression domains using cortically-biased kidney scRNA-Seq data from^26^. Only the hematopoietic cluster was omitted. (C) Genome browser view of the chr17 fragment contained in the *Six2TGC* BAC with alignment of ATAC-seq tracks and differential Hi-C called loops for *Six2TGC* (TGC)-containing NPCs vs. control (CON). Loop #1; *Six2* and *Six3* promoter interactions. Loop #2; the *Six2* distal enhancer and *Six3* promoter. Loop #3; junctional interactions of two co-integrated BAC (see Figure 2). (D) Juicebox visualization of the chr17 segment shown in (C) with *Six2TGC* on the x-axis and control on the y-axis to visualize differential contacts and position of the TAD boundary. Dashed blue circles emphasize the absence of the indicated interactions in the control condition. (E) 3C-PCR product confirming physical interaction suggested by Loop #2, absent from the control library. ‘M’ denotes DNA size marker.

To ask if reduction in nephrons was caused by local disruption at the integration site or by aberrant regulatory interactions within the transgene possibly driving *Six3* expression, we generated Hi-C libraries from freshly isolated NPCs from P0 *Six2TGC^+/tg^*, *Six2TGC^+/tg^; Tsc1^+/f^* and a control *Cited1^CreERT2^* transgenic line (herein *Cited1^+/GCE^*) representing mouse lines with low, restored, and control nephron numbers, respectively. Previous studies established a conserved topologically associating domain (TAD) boundary insulating the *Six3* promoter from engaging with *Six2*-DE^19^ within chr17^20^. ATAC-seq from P2 *Six2TGC^+/tg^*; *Tsc1^+/flox^* NPCs confirmed that the *Six3* promoter was accessible in BAC-containing NPCs but not on chr17 in controls, consistent with transgene-specific interaction (Figure 1C). Further, differential loop calling using HiCCUPS^21^ identified several unique contacts in *Six2TGC^+/tg^* that are absent in the control condition (Figures 1C-D and Supplemental Table S2), including contacts between the *Six2*-DE and the *Six3* promoter. We validated this interaction by locus-specific chromatin conformation capture (3C)-PCR followed by sequencing on libraries generated from P0 nephrogenic zone cells isolated from *Six2TGC^+/tg^*and control mice (Figure 1E). Importantly, such products were absent from control NPCs, and were likewise not detected in control 4C libraries of *Six2*-DE interactions^20^. Collectively, the Hi-C, 3C, ATAC and RNA expression studies converge on the conclusion that the *Six2*-DE activates the *Six3* promoter within the transgene, which we hypothesize contributes to reduction in nephron numbers.

### The *Six2TGC* allele integrated within chromosome 1 and does not alter local transcription

To investigate whether the allele architecture and integration of the *Six2TGC* transgene might also contribute to reduced nephron number by disrupting endogenous gene regulation in NPCs, we submitted splenocytes from *Six2TGC^+/tg^*mice to Cergentis (Utrecht, Netherlands) for targeted locus amplification (TLA)^22^ to determine the transgene organization and integration site. TLA identified a candidate region for transgene integration in a 75kb span within a large intron of *Cntnap5a* located on chr1 (Supplemental Figure S1A). These results established that the transgene contained the BAC vector backbone (Supplemental Figure S1B), and that the integration breakpoint occurred within the body of the BAC. Since sequencing reads mapped to the mouse genome cannot differentiate the BAC from the endogenous sequence on chr17 we leveraged our Hi-C datasets to identify interactions between the transgene and neighboring genomic sequences^23, 24^ by aligning the genomes of two transgenic lines, *Six2TGC^+/tg^* and *Cited1^+/GCE^* (Supplemental Figure S2A). As expected, contact maps showed scarce interactions between chr1 and chr17 in the control (*Cited1^+/GCE^*) samples in full genome view (Supplemental Figure S2A) or between the two chromosomes in isolation (Supplemental Figure S1C). However, we could readily detect a hotspot of chr1/chr17 interaction in *Six2TGC^+/tg^* (Supplemental Figure S2A, blue circle, and Figure 2A) and a chrX/chr14 interaction in *Cited1^+/GCE^* mice (*Cited1* is located on the X chromosome; Supplemental Figure S2A, red circle). Both interactions were specific to the respective transgenic lines. At larger magnification, *Six2TGC^+/tg^* interaction frequency peaks near the putative integration site in *Cntnap5a* (Figure 2Aa) and *Cited1^+/GCE^*interaction frequency peaks near the putative integration site in *Usp54* (Supplemental Figure S2B,C). Note that unlike the contiguous chr17 signal on chr1, visualization of the data using Juicebox^21^ indicated a 450KB gap in the chr14 region with strongest interaction with the *Cited1* gene region on ChrX, located between *Usp54* and *Ap3m1* (Supplemental Figure S2B,C). Differential peak calls between *Six2TGC^+/tg^*-containing libraries and *Cited1^+/GCE^*overlap with the Hi-C gap, consistent with a large deletion near the *Cited1^CreERT2^*transgene integration site on chr14. Overlay of ATAC-seq data shows that in this region, *Six2TGC^+/tg^*; *Tsc1^+/flox^* has peaks twice as high as the corresponding ones in *Cited1^+/GCE^* (IGV track below the Hi-C map in Figure S2B, enlarged in Figure S2C). Conversely, differential peaks calls on the chrX region corresponding to the BAC used to construct the *Cited1^CreERT2^* transgene were four-fold higher in *Cited1^+/GCE^* than in *Six2TGC^+/tg^*, consistent with the report of four integrated BAC copies in this transgenic line^25^. While we have not characterized this line further, we anticipate that *Cited1^GCE/GCE^*animals will have phenotypic consequences due to the large deletion at the integration site.

**Figure 2.**
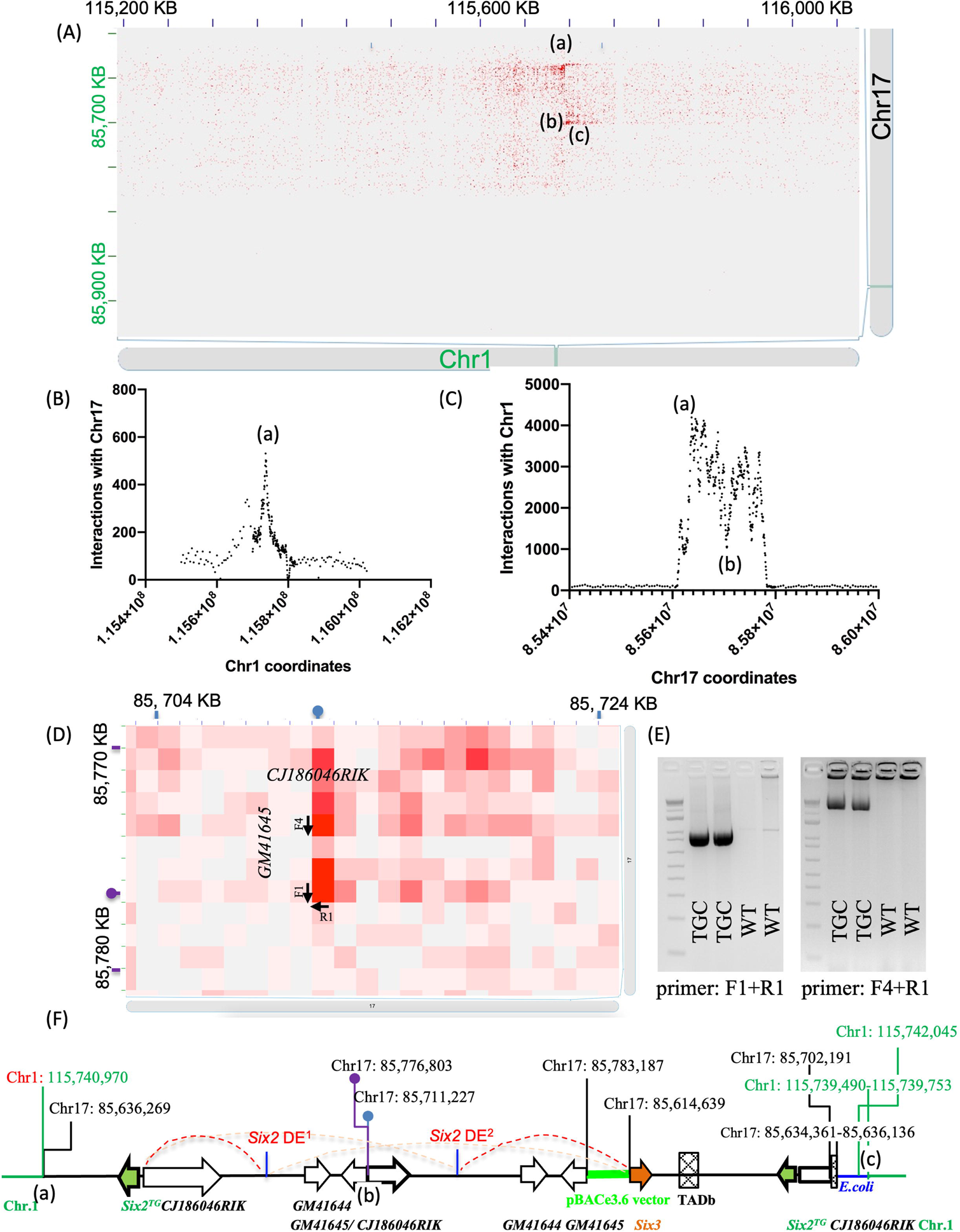
Hi-C analysis identifies the chromosomal integration site of *Six2TGC*. (A) Juicebox visualization of chr17-chr1 interchromosomal contacts in *Six2TGC* reveals the integration site. (B) Quantification of reads confidently mapping to chr1 with mate pairs mapping to the chr17 BAC sequence using the python module *pysam* is consistent with a single integration site, confirmed by PCR. (C) Quantification of chr17 reads with the mate pair mapping to chr1 reveals several candidate breakpoints within the BAC. (D-E) High-resolution view of the Hi-C data identifies the inter-BAC junction and 3’ boundary sequences, confirmed with the indicated PCR primers. ‘WT’ denotes wild-type control. (F) Combined analyses of Hi-C data validated by PCR product sequencing suggests the *Six2TGC* allele architecture proposed here, with the annotated boundaries and breakpoints listed. Green line denotes the pBACe3.6 vector; *Six2* DE denotes the *Six2* distal enhancer; *E. coli* denotes contaminating, co-integrating bacterial genomic DNA; green arrow denotes the eGFP-Cre transgene; orange arrow denotes the *Six3* locus. Schematic not to scale.

To leverage the HiC data to define the *Six2TGC* integration site precisely we used the python module *pysam* to implement a sliding 5kb-window across the *Cntnap5a* gene region and quantify high-quality chr1 reads with mate pairs mapping confidently to chr17. This method identified a single peak within a 5kb genomic window, mapping to the region predicted by TLA, containing the highest number of interactions with the BAC sequence (Figure 2B). PCR using genomic DNA isolated from deceased, presumptive *Six2TGC^tg/tg^* homozygotes at P0 confirmed the integration site on chr1. PCR screening of candidate BAC breakpoints from this analysis amplified the 5’ chr1-transgene junction, confirmed by sequencing (Supplemental Figure S1D).

Having mapped the transgene integration site on chr1, we analyzed ATAC and RNA-seq^26^ data for changes in transcript levels of genes near the integration site, as we did for chr14 in *Cited1^+/CGE^*. No differential transcripts, ATAC peaks, or novel/disrupted Hi-C loops were detected within a 3MB region centered at the transgene integration site (Supplemental Figure S1E). No change in long range interactions within 30MB region was detected (Supplemental Figure S1F), indicating that the transgene integration did not disrupt expression of any endogenous gene on chr1 that might contribute to the low nephron phenotype. Furthermore, no interactions between the *Six3* promoter and chr1 were detected by *pysam* scanning.

### SIX3 interferes with SIX2 function and disrupting *Six3* within the *Six2TGC* allele restores nephron endowment

To test whether *Six3* misexpression was detrimental to nephron endowment in *Six2TGC^+/tg^* mice, we further characterized the transgene structure to inform the design of a potential rescue strategy. We detected two copies of *Cre*^27, 28^ suggesting that two partial or complete BACs integrated in tandem with an unknown orientation. Therefore, we implemented a similar sliding window approach as above, scanning across the BAC sequence and counting interaction pairs of chr17 with chr1. Closer inspection of the interactions between chr1 vs. chr17 using Juicebox^21^ suggested that the 3’ boundary of the transgene may lie in the middle of the BAC (Figures 2Ab,c, C). We then determined the exact sequence at the transgene-chr1 3’ junction by inverse PCR using primers specific for the 3’ side chr1 sequences. Sequencing the *Six2TGC*-specific PCR products revealed a complex integration junction containing small, duplicated fragments from both chr17 and chr1 as well as a 635-bp fragment of *E. coli* genomic DNA (Supplemental Figure S1G). Examination of the Hi-C data at 1kb window (Figure 2D) predicted an inter-BAC junction between two copies of chr17 sequences, corresponding to one of the novel Hi-C loops found in the *Six2TGC^+/tg^*samples (loop #3 in Figure 1C). Following PCR amplification, we sequence-verified the 5’ and 3’ junctions of the transgene integration site, as well as the junction between the two copies of the transgene (Figures 2E-F and Supplemental Figure S1H). The entire *Six2TGC* allele contains two incomplete copies of the original BAC, with two *Six2*-DE, one copy of the *Six3* gene containing both coding exons and its proximal promoter, and contaminating bacterial sequences (vector backbone and genomic *E. coli* DNA) which may also explain variegation and mosaicism in Cre activity as reported by users^29^. This architecture and all junctional coordinates (mm10) are annotated in Figure 2F. Note that the loops reported in Figure 1C cannot be assigned in Figure 2F to a particular *Six2-DE* enhancer in a custom *Six2TGC* genome because duplicated regions generate indistinguishable read pairs.

To test the hypothesis that aberrant *Six3* expression disrupts kidney development in *Six2TGC^+/tg^* mice and consequently endeavor to correct this deficit, we pursued CRISPR-mediated editing. We used a single gRNA (target sequence GGAGAACCCCTACAGTTCTG <u>GGG</u>) targeting the intron of *Six3*. Twelve CRISPR-targeted *Six2TGC* founder mice were bred to wild-type CD1 mice. To screen for specific disruption of the *Six3* gene in the transgene and not the endogenous *Six3* on chr17, kidneys were isolated from the resultant F1 offspring at P0 and total RNA was extracted from nephrogenic zone cells followed by RT-PCR detection of transgene-derived *Six3* transcripts spanning exon 1 and the exon 1/exon 2 junction. We identified two founder mice transmitting a modified *Six2TGC* allele that failed to produce the *Six3* transcript containing sequences from exon 2 (Figure 3A). Founder #77 was selected for further breeding and comprehensive analysis, with SNP genotyping that distinguished the B6-derived *Six3* locus in the *Six2TGC* allele from the endogenous (CD1-derived) chr17 locus showing complete deletion of the *Six2TGC*-associated *Six3* exon 2 (Figure 3B, Supplemental Figure S1I). Importantly, nephron numbers in *Six2TGC*^Δ*Six3CT/+*^ mice lacking the *Six3* exon2 region within the transgene were restored to that of non-transgenic littermate controls (Figure 3C). When bred to homozygosity, *Six2TGC*^Δ*Six3CT/*Δ*Six3CT*^ kidneys were only slightly reduced in size compared to *Six2TGC*^Δ*Six3CT/+*^ samples and developmentally appropriate compared to the aplastic *Six2TGC^tg/tg^*kidneys (Figure 3D). Together, these results confirm that ectopic expression of *Six3* from the transgene is responsible for disrupting the establishment of nephron number in the *Six2TGC^+/tg^* mice. The mild hypoplasia of *Six2TGC*^Δ*Six3CT/*Δ*Six3CT*^ kidneys may result from the expression of a truncated SIX3 protein. Homozygous viability was not rescued due to a severe cleft palate which may be caused by regions of *Six3* that were not disrupted in *Six2TGC*^Δ*Six3CT*^ (Supplemental Figure S1J)

**Figure 3.**
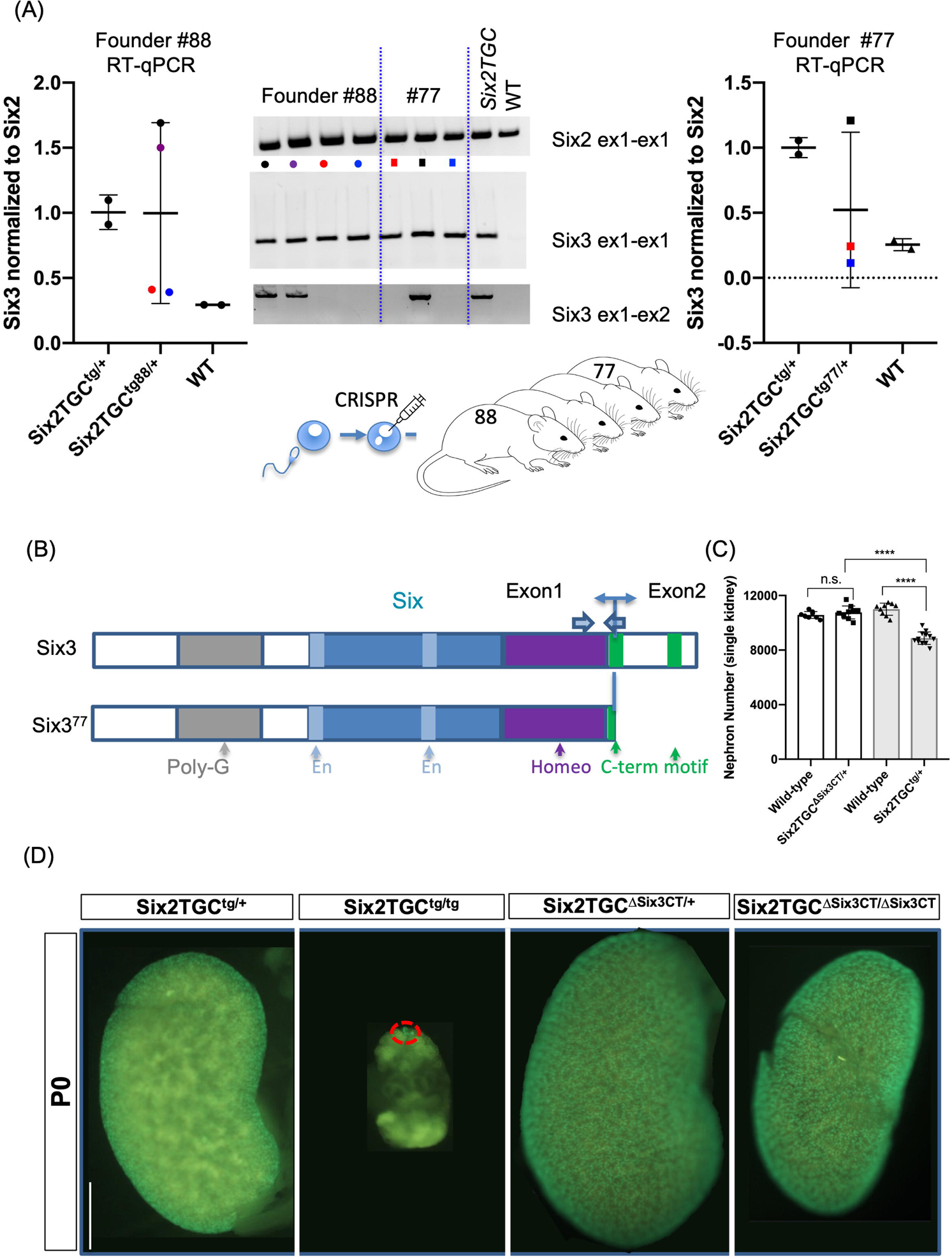
CRISPR-mediated disruption of *Six3* within the BAC restores nephron number. (A) CRISPR and RT-qPCR screening strategy identifies founders (#77 and # 88) with loss of *Six3* exon 1-2 splicing; *Six2* and *Six3* exon 1 PCR products serve as positive controls for mRNA isolation from NPCs and transgene presence, respectively. (B) Predicted protein product domains for modified transgene *Six3* allele (*Six2TGC*^Δ*Six3CT*^) compared to endogenous *Six3*. (C) Nephron number estimated by acid maceration at ≥P28 shows normalized nephron numbers in *Six2TGC*^Δ*Six3CT*^ mouse as compared to the original *Six2TGC* transgenic line. One-way ANOVA with Tukey’s multiple comparisons test was performed in GraphPad Prism version 8 to evaluate statistical significance, ‘n.s.’ denotes not significant, **** denotes p ≤ 0.0001. (D) Surface GFP visualization of P0 kidneys isolated from the indicated genotypes; scale bar denotes 1mm.

### Possible mechanisms underlying deleterious activity of SIX3

Whereas *Six2TGC^tg/+^* mice have reduced nephron number, *Six2TGC^tg/tg^* mice are born with severely hypoplastic kidneys, a phenotype similar to that seen in the *Six2^-/-^* mice^30^ (Figure 3D). If both partial BAC copies are to be counted as a single unit, the *Six2TGC^+/tg^* mouse has a BAC/chr17 ratio of 1:2 and a *Six3^CT^/Six2* ratio of 1:2. The *Six2TGC^tg/tg^* homozygous mice have a BAC/chr17 ratio of 2:2, and a *Six3^CT^/Six2* ratio of 2:2; the *Six2TGC*^Δ*Six3CT/*Δ*Six3CT*^ mice have the same BAC/chr17 ratio of 2:2, but a *Six3^CT^/Six2* ratio of 0:2. To determine whether the *Six3^CT^/Six2* ratio was the main determinant of *Six2TGC*-mediated interference with nephrogenesis, we analyzed *Six2TGC^+/tg^*;*Six2^+/CE^* animals with the same BAC/chr17 ratio as the *Six2TGC^+/tg^*transgene (1:2) but a *Six3^CT^/Six2* ratio of 1:1, equal to that of *Six2TGC^tg/tg^* (2:2 *Six3^CT^/Six2*). Whole kidneys from these animals at E17.5 and/or P0 were examined via surface visualization of GFP, expressed in the peripheral NPCs (Figure 3D and Supplemental Figure S3). While nephron number is increased in *Six2* heterozygotes^12^, NPCs were extinguished by E17.5 in *Six2TGC^+/^*^tg^;*Six2^+CE^* kidneys (Supplemental Figure S3). Collectively, these data suggest that the stoichiometry of *Six3^CT^/Six2*, and not the absolute BAC copy number or intact chr1, is the critical determinant of NPC survival and nephron endowment. Accordingly, *Six2TGC^tg/^*^Δ*Six3CT*^ kidneys at birth are much larger than *Six2TGC^tg/tg^* but smaller than *Six2TGC*^Δ*Six3CT/*Δ*Six3CT*^ which approximate normal size (Figure 3D and Supplemental Figure S3). Collectively, these analyses indicate that the negative impact of the *Six2TGC* maps to the C-terminal region of *Six3*.

The possibility of homodimerization of SIX family TFs, comprised of distinct subfamilies of closely related pairs *Six1/2, Six4/5,* and *Six3/6*^31^, has been discussed^32^, and heterodimerization was predicted by the AI program AlphaFold^33^ (Figure 4B). Since we could not detect SIX3 protein by IF with commercially available antibodies in control tissue, we co-transfected the physiologically relevant mouse metanephros-derived mk4 cells with tagged murine *Six2-GFP* and *Six3-FLAG* constructs to test whether SIX2 and SIX3 proteins are able to interact. Co-immunoprecipitation with the FLAG antibody (but not with isotype control) followed by western blot detection of SIX2 reproducibly detected direct interaction between SIX3-FLAG and SIX2-GFP (Figure 4C, showing one of three repeats). However, the reciprocal CoIP with anti-GFP antibody did not recover SIX3-FLAG, and this mechanism remains speculative.

**Figure 4.**
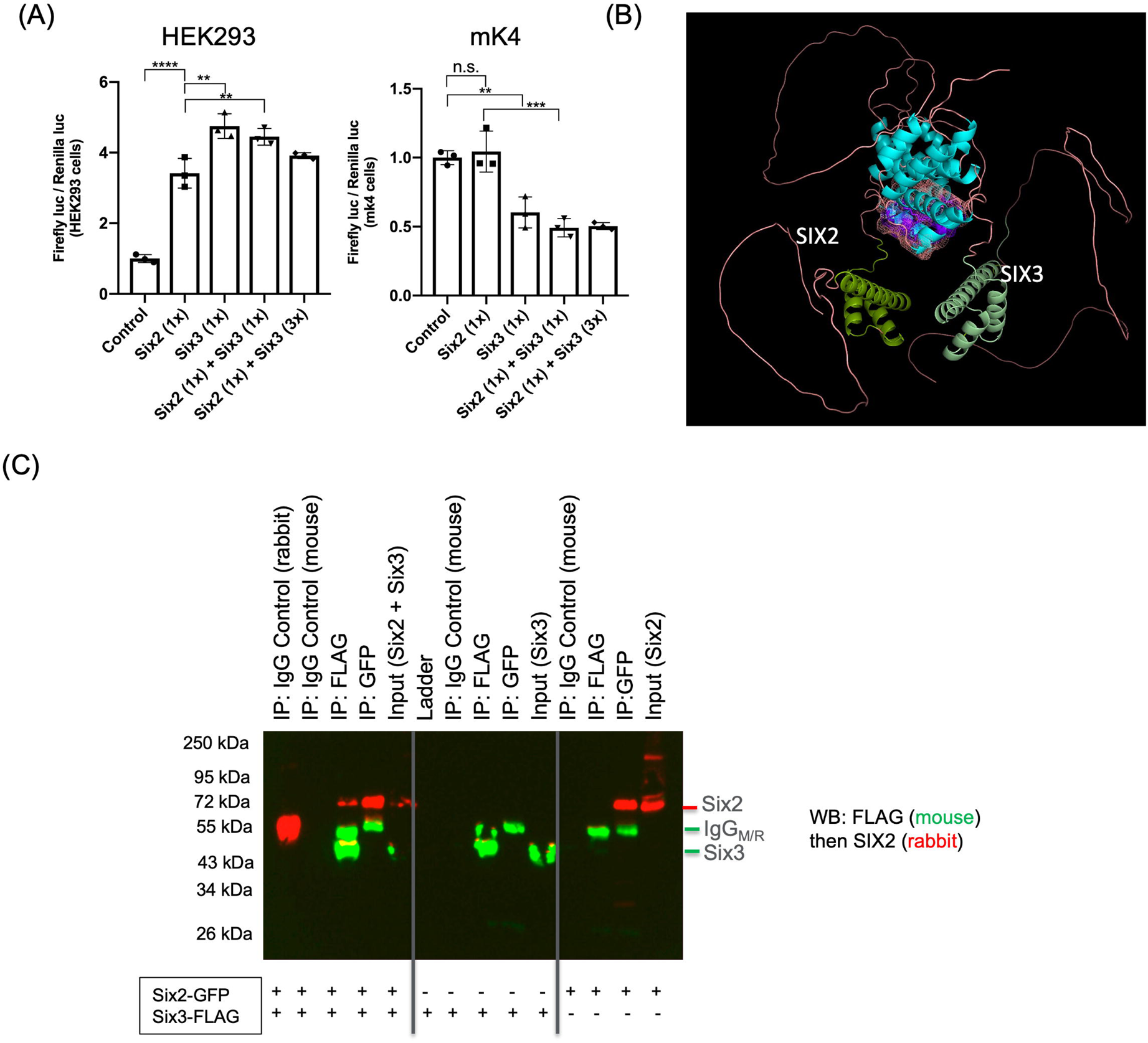
SIX3 interference with SIX2 activity. (A) *Six2* promoter luciferase reporter assay in human HEK293 and murine mk4 cells with varying ratios of *Six2* and *Six3* co-transfected. One-way ANOVA with Tukey’s multiple comparisons test was performed in GraphPad Prism version 8 to evaluate statistical significance, ‘n.s.’ denotes not significant, ** denotes p ≤ 0.01, *** denotes p < 0.001, **** denotes p ≤ 0.0001. (B) AlphaFold prediction of murine SIX2/SIX3 heterodimer structure. (C) Immunoprecipitation of FLAG-tagged SIX3 followed by Western blot for GFP-tagged SIX2 identifies physical association between SIX2 and SIX3 in co-transfected mk4 cells. ‘IgG’ denotes isotype control.

We further investigated whether SIX3 might interfere with SIX2-mediated transcriptional regulation. We first tested the impact of SIX2 and SIX3 overexpression on a *Six2* promoter-based luciferase reporter activated by *Six2* in HEK293 cells^34^. Both SIX2 and SIX3 activated the reporter in human HEK293 cells. In the mK4 cell line^35^, SIX2 failed to activate its own promoter or the reporter construct above baseline. By contrast, SIX3 modestly reduced reporter baseline activity even in the presence of SIX2 (Figure 4A), in accord with reports that SIX3 and SIX2 can act as activators or repressors^4, 36, 37^.

Although heterodimerization cannot be demonstrated *in-vivo* in the absence of a working SIX3 antibody, these data, together with the genetic dose-dependent interactions described above, support a scenario in which SIX3 produced from the *Six2TGC* transgene interferes with SIX2 activity in the regulation of NPC renewal. Combined, these observations are most consistent with a model in which the C-terminal domain of SIX3 protein plays a crucial role in its interference with SIX2 function in nephron development.

## DISCUSSION

Increased chromatin accessibility at putative SIX2 binding sites is characteristic of younger NPCs, consistent with the observation that SIX2 extinction is associated with differentiation^30^ in part by enabling canonical WNT activation at a subset of loci^4^. One possible explanation for how ectopic *Six3* expression reduced nephron number in *Six2TGC^+/tg^* mice, perhaps the most widely used Cre driver mouse for tissue-specific genetic studies of kidney development, is that the presence of a SIX3 protein containing its C-terminal domain interferes with the ability of SIX2 to antagonize WNT signaling. If this conjuncture is proven correct, all genetic models reducing or expanding nephron numbers analyzed thus far converge on the balance between FGF and WNT signal strength as key determinants of nephron endowment.

To arrive at this point we characterized the integration site and architecture of the *Six2TGC* allele and probed the mechanism that permits inappropriate expression of *Six3* in *Six2TGC*-containing NPCs. Other than *Six3* activation, no evidence of altered regulatory interactions or transcription within 50MB region centered on the integration site were detected. The *Six3* transcripts were isolated from the polysome in similar abundance to *Six2*, which together with our data showing CRISPR deletion of the *Six3*-exon2 in the transgene rescued nephron numbers indicate misexpressed SIX3 protein interferes with SIX2-mediated regulation of nephrogenesis. We demonstrate that physical association of SIX3 with SIX2 is possible, and that the BAC/chr17 (presumably, the SIX3/SIX2) ratio in NPCs is detrimental to kidney development and nephron numbers. It has been reported that the *brachyrrhine* (*Br*) mutation causes reduction in *Six2* expression and concomitant ectopic *Six3* expression in NPCs due to inversion of the *Six2* and *Six3* genes relative to their distal enhancers within the locus^20^. In contrast to *Six2^+/-^*mice that display increased nephron number, the *Br/+* mice exhibited reduced nephron number and chronic kidney failure^38^. However, despite the demonstration of ectopic *Six3* expression in NPCs activated by *Six2* enhancers in the *Br/+* kidneys, a role for SIX3 in the *Br/+* renal defects has not been considered. We leveraged these observations to test the hypothesis that SIX3 misexpression in the kidney is deleterious by generating a modified transgene which disrupts the C-terminus of *Six3* and demonstrated that this minimal change was sufficient to rescue nephron numbers and abrogate the negative effect of *BAC* dosage.

Our studies of the *Six2TGC and Cited1GCE* transgenes provided a roadmap toward rapid mapping of classical, commonly used transgenes. Transgenic mouse lines, including *Cre* lines mediating conditional loss of gene expression^39^, are powerful tools that enabled decades of genetic and molecular studies of development and disease in targeted animals and cells. Random genomic integration of a single BAC or a tandem array confounds interpretation of results due to disruption of endogenous gene expression and/or induction of large deletions near the integration site^40^ as has occurred in *Cited1^GCE/+^* mice. Several experimental approaches have been applied to identify transgene integration sites, including TLA^22^ and more recently nanopore sequencing^41, 42^. Herein, we demonstrate how inter-chromosomal interactions in Hi-C datasets, coupled with transgene specific ATAC data, can rapidly identify transgene integration sites, provide an estimate of copy number, and elucidate BAC and locus architecture post-integration. Importantly, our straightforward approach will facilitate evaluation of unintended transcriptional dysregulation arising from a transgene or near the integration site, and the presence of modifiers due to haploinsufficiency.

While many genetic and environmental factors can impact nephron number^10^, the list is currently incomplete. While this modified transgene will provide an improved reagent for investigating NPCs without significant disruption to normal nephron formation, a minimally disruptive redesign of the transgene will be required to ensure viability. This effort is underway. The datasets presented herein include high resolution Hi-C libraries that complement a rich resource of chromatin conformation and accessibility data, aiding ongoing and future investigations of nephron development and number determination. Moreover, ATAC and Hi-C data from multiple developmental ages and strains will help elucidate regulatory mechanisms underlying variability between strains, developmental timepoints, and mutants with downstream clinical implications for identifying interventions in populations vulnerable to early onset CKD secondary to low nephron endowment.

## Methods

### Animals

The following mouse lines were used in this study: *Tg (Six2-EGFP/cre)1^Amc^* (Jax Stock No: 009600; herein *Six2TGC*), *Six2-<u>C</u>re<u>E</u>RT2*, (herein *Six2^+/CE^*), *Tsc1^f/f^*, and *Cited1-CreERT2-GFP^+/tg^* (herein *Cited1^+/GCE^*). These lines were maintained by breeding to CD1 mice (Charles River, strain code: 022). In mouse experiments using embryos, embryonic day (E) 0.5 was designated as noon on the day a mating plug was observed. All mice were maintained in the Cincinnati Children’s Hospital Medical Center (CCHMC) animal facility according to the animal care regulations. Our experimental protocols (IACUC2018-0107/0108 and IACUC2021-0067/0077) are approved by the Institutional Animal Care and Use Committee of CCHMC. Animals were housed in a controlled environment with a 14h light/10h dark cycle, with free access to water and a standard chow diet. Genotyping was performed on DNA isolated from the toe or tail using the primer sets as follows:

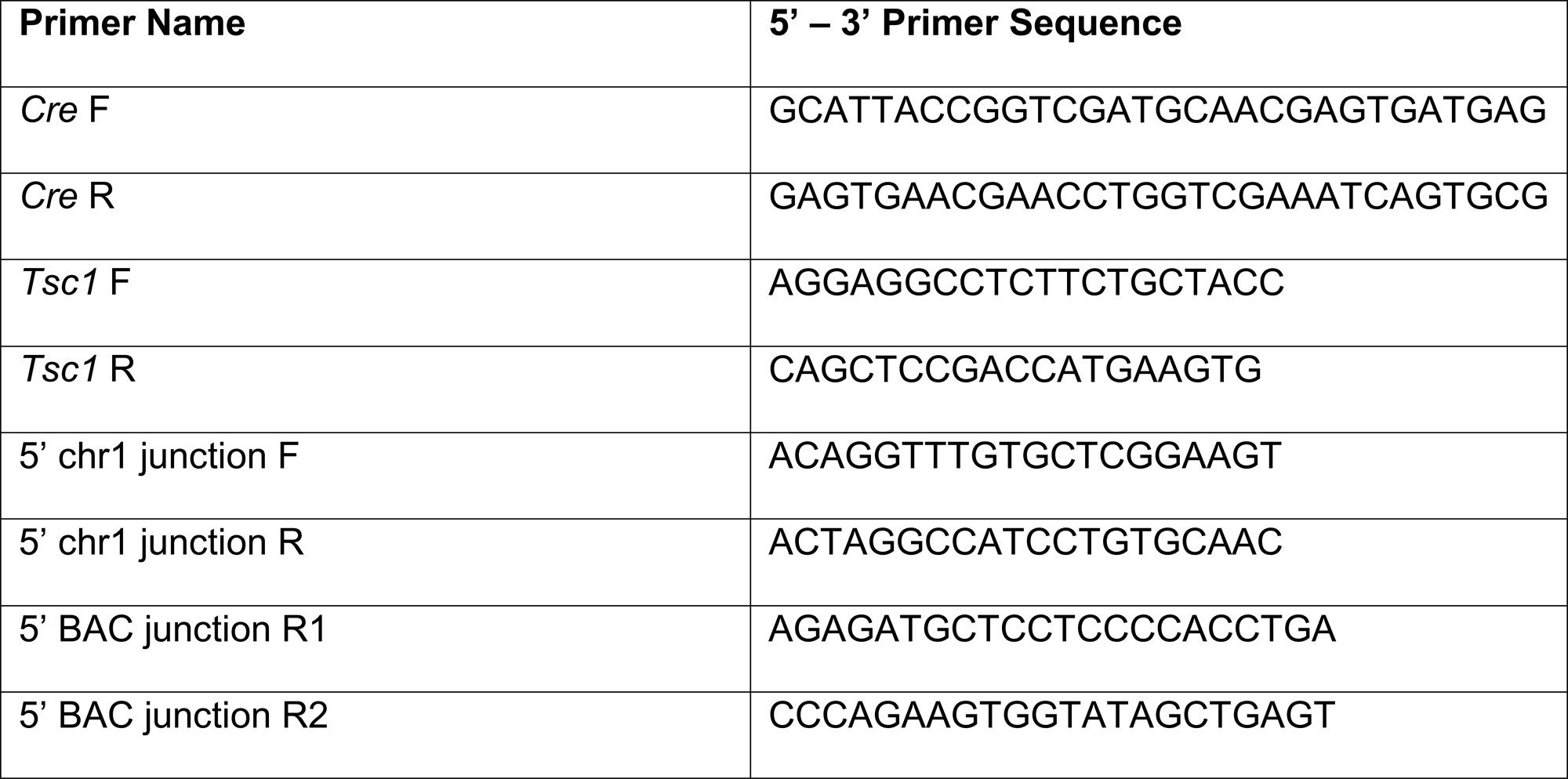

### ATAC-seq

For sample library preparation, we followed the Omni-ATAC method outlined by^43, 44^ and purified Tn5 was generated as described^45^. Briefly, 50,000 nuclei from FACS-sorted NPCs (isolated as described in^26^) were processed for Tn5 transposase-mediated tagmentation and adaptor incorporation at sites of accessible chromatin. NPCs were pelleted and washed with ice-cold PBS. The pellet was resuspended in ATAC-Resuspension Buffer (10 mM Tris-HCl [pH 7.4], 10 mM NaCl, 3 mM MgCl_2_ 0.1% NP40, 0.1% Tween-20, 0.01% Digitonin) and incubated on ice for 3 minutes. The lysed cells were washed in ATAC-Wash Buffer (10 mM Tris-HCl [pH 7.4], 10 mM NaCl, 3 mM MgCl_2_, 0.1% Tween-20), inverted 3 times, and the nuclei pelleted. The nuclei were resuspended in ATAC transposition mix (10 mM Tris-HCl [pH 7.6], 10 mM MgCl_2_, 20% Formamide, 100 nM Tn5 transposase) and incubated at 37°C for 30 minutes in a thermomixer at 1,000 RPM. Following tagmentation, the DNA fragments were purified using a Zymo DNA Clean and Concentrator Kit, and library amplification was performed using customized Nextera PCR primer Ad1 in combination with any of Ad2.1 through Ad2.12 barcoded primers as described. The quality of the purified DNA library was evaluated utilizing an Agilent Bioanalyzer 2100 using High Sensitivity DNA Chips (Agilent Technologies Inc.). The samples were pooled at a concentration of 5 nM and run on an Illumina HI-SEQ 2500 sequencer (Illumina) to obtain paired-end reads of 75 bases (PE75). Each ATAC sequencing condition was performed in biological triplicate. Sequences were aligned to the mm10 genome. CSBB-v3.0. pipeline with embedded fastqc, bowtie2 and macs2 was used for quality check, mapping, and peak calling. Duplicate mapped reads were removed before peak calling. Motif enrichment analysis was performed using HOMER^46^.

### 3C

Nephrogenic zone cells were isolated from *Six2TGC^+/tg^*and control littermate mice at P0. Four to six kidneys per sample were digested with 2.5 mg/ml collagenase D and incubated at 37°C for ∼20-25 minutes in a Thermomixer R at 1400RPM. 3C was performed as described in^47^. Approximately 10 million cells were pooled per replicate and *DpnII* was used as the restriction enzyme. Primers used to detect the *Six3* promoter and *Six2*-DE interaction are as follows, 5’-3’: CCCTTACGTCCTTCCTCCTC and CTATGGTGATTGGAAACCCG.

### Hi-C sample preparation

The Arima Genomics low input Hi-C protocol was performed according to manufacturer instructions. Briefly, P0 *Six2TGC^+/tg^,* P0 *Six2TGC^+/tg^; Tsc1 ^+/f^* and P0 *Cited1^+/GCE^*nephron progenitor cells were isolated by FACS for GFP+ cells as follows: kidneys were dissected in ice-cold PBS and the capsule was removed. Four to six kidneys per sample were digested with 2.5 mg/ml collagenase D and incubated at 37°C for ∼20-25 minutes in a Thermomixer R at 1400RPM. Kidneys were examined under a fluorescent microscope to ensure removal of nephron progenitors (GFP+). The kidneys were removed, and the cells were washed twice in 0.1% BSA/PBS. Cells were isolated using a Sony SH800S instrument and counted with a hemocytometer. Approximately 800,000 cells per genotype were pooled, with 3 biological replicates generated per genotype. Libraries were prepared using the Swift Biosciences ACCEL-NGS® 2S Plus DNA Library Kit, individually barcoded, and pooled into a single sample. The pooled library was submitted to Novogene for sequencing via Illumina NovaSeq 6000. Hi-C library sequencing reads were processed with Juicer (v1.5.7)^21, 48^. For each of the three biological conditions, the pooled triplicates yielded ∼1 billion (B) total sequenced read pairs and ∼330 million (M) intra-chromosomal long-range (>20 kilobases (kb)) fragments.

### Identification of BAC integrations by Hi-C

BAM files utilized for BAC integration analysis were initially mapped to the mm10 reference genome by bwa (version: 0.7.17-r1188) with parameters “bwa mem-SP5M” and samblaster (v0.1.25) to mark the PCR duplicated reads. Only high-quality reads (MAPQ>30 with both ends uniquely mapped and no PCR duplication) were utilized for the analysis. Customized python script (available in Github.org) with default parameters was utilized to estimate the number of high quality paired-end reads with one end mapped in chr17 and the other end mapped in the targeted regions in chr1. All the analyses ware repeated across three biological replicates in each condition, and the results were consistent. The results of the merged bam files from three biological replicates were utilized for the figures. Merged bam files in each condition were utilized to get more reads in small windows. Hi-C loop analysis was performed using Juicer (v1.5.7)^21, 48^, with a resolution of 10kb in HiCCUPS.

### RNA Isolation and RT-qPCR

RNA from cells was extracted using the PureLink RNA Mini kit (Invitrogen) and cDNA was synthesized with SuperScript II reverse transcriptase (Invitrogen) following the manufacturer’s instruction. Quantitative PCRs were performed using iTaq Universal SYBR Green Supermix (Bio-Rad) on the StepOnePlus RT PCR system (ThermoFisher). Data were analyzed using the Delta-Delta-CT method. Primers used are as follows, listed from 5’ – 3’: *Six2* F ‘GCAAGTCAGCAACTGGTTCA’, *Six2* R ‘CTTCTCATCCTCGGAACTGC’, *Six3* F ‘AGAACAGGCTCCAGCATCAG’, *Six3* R ‘TACCGAGAGGATCGAAGTGC’, *Six3* exon1 F ‘CCGGAAGAGTTGTCCATGTTC’, *Six3* exon1 R ‘CGACTCGTGTTTGTTGATGGC’.

### Surface imaging

E17.5 and P0 kidneys were dissected in cold PBS and GFP was visualized using a Lecia stereo-fluorescent microscope.

### Co-immunoprecipitation

24 hours prior to transfection, mk4 cells were seeded at a density of 250,000 cells per well on a 6-well plate. Murine tagged *Six2* (OriGene CAT# MG20410) and *Six3* (OriGene CAT# MR204911) constructs were co-transfected in mK4 cells at a 1:1 ratio in Opti-MEM (Gibco). DNA (1 µg per well) and Lipofectamine 2000 (Invitrogen) were incubated for 20 minutes at room temperature prior to transfection. 48 hours after transfection, cells were collected by scraping for protein isolation in CoIP Lysis Buffer (50 mM Tris pH 7.4, 150 mM NaCl, 1 mM EDTA, 1% Triton X-100) with protease inhibitors. Co-immunoprecipitation was performed per manufacturer’s instructions for the Pierce^TM^ Protein A/G Magnetic Beads (Thermo Scientific) kit, and samples were eluted into 2X sample buffer and heated at 65 degrees for 10 minutes for subsequent detection by western blotting. Antibodies used are as follows: for immunoprecipitation (OriGene, CAT #TA-50011-3 and Origene, #TA150041); for western blot: anti-Six2 (1:500, 11562-1-AP, Proteintech) and (OriGene, CAT #TA-50011-3, 1:2000) with donkey anti-rabbit HRP and sheep anti-mouse HRP secondaries (1:10000).

### Luciferase assay

The tagged murine *Six2* and *Six3* constructs indicated above, and reporter constructs from^34^ were co-transfected into HEK293 and mk4 cells plated at 50,000 cells/well on 96-well plates in triplicate; 100ng DNA per well. Luciferase reporter assays were performed as described in^34^, and according to the manufacturer’s instructions (Dual-Glo Luciferase Assay System, Promega). One-way ANOVA with Tukey’s multiple comparisons tests were performed in GraphPad Prism version 8 to evaluate statistical significance.

### Nephron counts

HCl maceration of whole kidneys was performed according to^49, 50^. Kidneys were isolated from ≥ P28 animals, the capsule removed, minced using a razor blade and incubated in 6N HCL for 90 min. The dissociated kidneys were vigorously pipetted every 20-30 minutes to further disrupt the kidneys. After incubation, 5 volumes of distilled water were added to the samples followed by incubation at 4°C overnight. 100µl of the macerate was then pipetted into a cell culture dish and glomeruli were counted in triplicate (three aliquots) for each sample. A single experimenter, blinded to the genotypes of kidneys being scored, performed all counts. Genotyping was performed after the counts on DNA isolated from the liver, spleen or tail using the primer sets reported above. One-way ANOVA with Tukey’s multiple comparisons tests were performed in GraphPad Prism version 8 to evaluate statistical significance. Both kidneys were counted for each animal with males and females represented.

### Generation of mice with Six3 disruption in the BAC

We targeted *Six3* intron with a single guide RNA (target sequence: GGAGAACCCCTACAGTTCTG <u>GGG</u>) according to the optimal on- and off-target scores from the web tool CRISPOR (http://crispor.tefor.net ^51^) and the ideal location (e.g., low conservation and distant from the splice sites). To form ribonucleoprotein complexes (RNPs), 60 ng/ul synthetic single-guide RNA (IDT) and 80 ng/ul Cas9 protein (IDT) in Opti-MEM (Thermo Fischer) were incubated at 37°C for 15 min. The zygotes from super-ovulated *Six2TGC* female mice on the CD1 genetic background were electroporated with 7.5µl RNPs on ice using a Genome Editor electroporator (BEX Co. Ltd); 30V, 1ms width, and 5 pulses with 1s interval. Two minutes after electroporation, zygotes were moved into 500µl cold M2 medium (Sigma-Aldrich), warmed to room temperature, and then transferred into the oviductal ampulla of pseudo-pregnant CD1 females. 31 F0 CRISPR pups were born, of which 12 contained the *TGC* allele based on Cre PCR. We screened for *Six3* expression in the F1 offspring kidneys as described in the main text.

### Availability of data and code

The raw fastq and processed files are made available in the GEO database via the following accession codes for Hi-C sequencing data (https://www.ncbi.nlm.nih.gov/geo/query/acc.cgi?acc=GSE222062), and ATAC-Seq (https://www.ncbi.nlm.nih.gov/geo/query/acc.cgi?acc=GSE224324). The python code utilized to characterize the BAC integration sites (Figures 2B-C) is accessible via GitHub (https://github.com/AJPerl/Transgene-Integration-Site) (for Figures 2B-F).

## ACKNOWLEDGMENTS

We thank the Transgenic Animal and Genome Editing Core staff, as well as the Research Flow Cytometry Core (supported by NIH S10OD023410) at Cincinnati Children’s Hospital Medical Center (CCHMC). We also acknowledge funding support from the National Institute of Diabetes and Digestive and Kidney Diseases (NIH R01 DK106225 to R.K.; F30 DK123841 to A.J.P.).

## AUTHOR CONTRIBUTIONS

R.K. and A.J.P conceived the project; R.K., A.J.P., R.J., H.L., Y.H., M.H., and N.A. designed and/or conducted experiments and interpreted data; Y.L., A.J.P., P.C. and R.K. performed bioinformatic analysis of ATAC-seq and Hi-C data. R.K. and A.J.P wrote the manuscript with input from all authors.

