## Supplemental Information for "Reduced nephron endowment in the common *Six2-TGC^tg^* mouse line is due to *Six3* misexpression by aberrant enhancer-promoter interactions in the transgene"

### Supplemental Figure S1

(A)

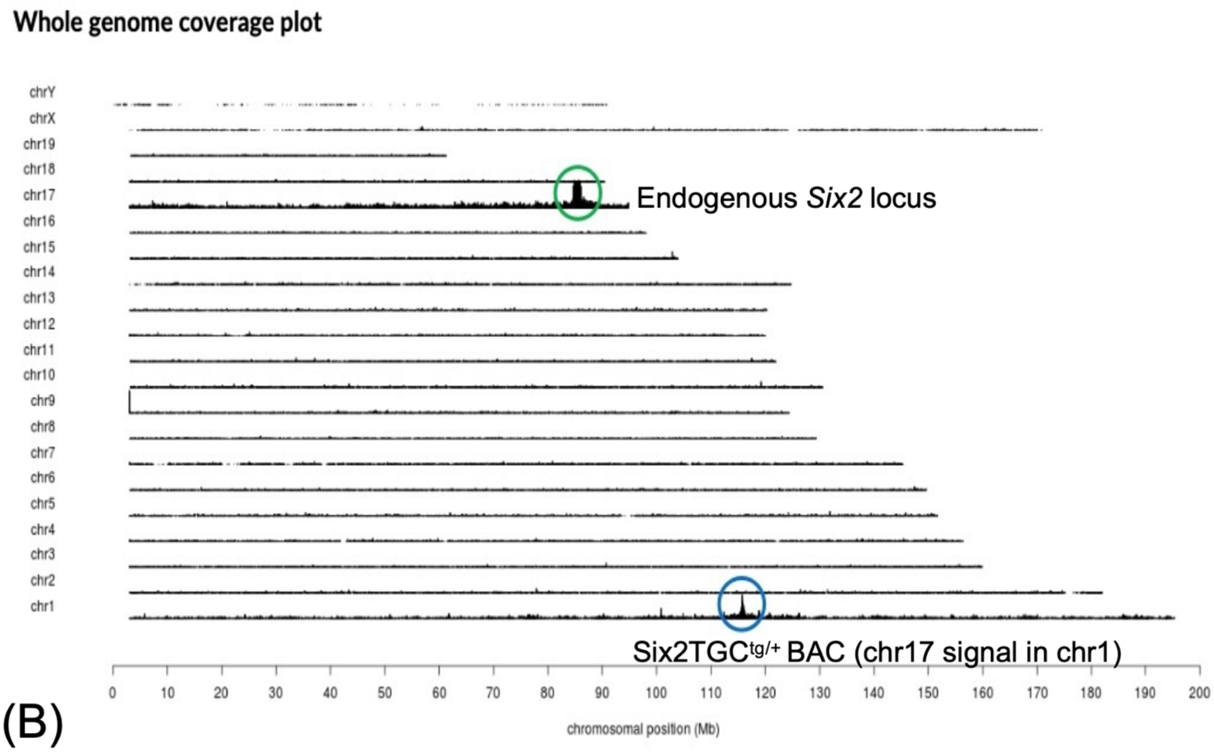


(B)

Good coverage is observed across the pBACe3.6 sequence (backbone) within pBACe3.6: 2,800-11,612 (flanked by the BamHI sites used to generate the BAC library). At the edges of the coverage, the fusions to the mouse BAC sequences are found:

Chr17:85,614,633 (head) fused to pBACe3.6:2,800 (head)
GCTTTTGTGTGTCCACCTCGAGCAGATCCGGAGCTGGCCGACCCCAGACTGATTCTCAACAGGTGGCTGGGTAAGT
CTGTTTAAAGAATTCCGCGGATCCTTCTATAGTGTCACCTAAATGTCGACGGCCAGGCGGCCGCCAGGCCTACCC

pBACe3.6:11,612 (tail) fused to chr17:85,783,185 (tail) with 8 bp inserted
TTGTAGGACTATATTGCTCTAATAAATTTGCGGCCGCTAATACGACTCACTATAGGGAGAGGATCCGCGGAATTCTT
GGTTCCATAACAAGTCCACTTTACTGTTTTCTTCGATTGGCTCCAAGAAGCACAAGTTCTTAGATGGAGGGTGC

(C)

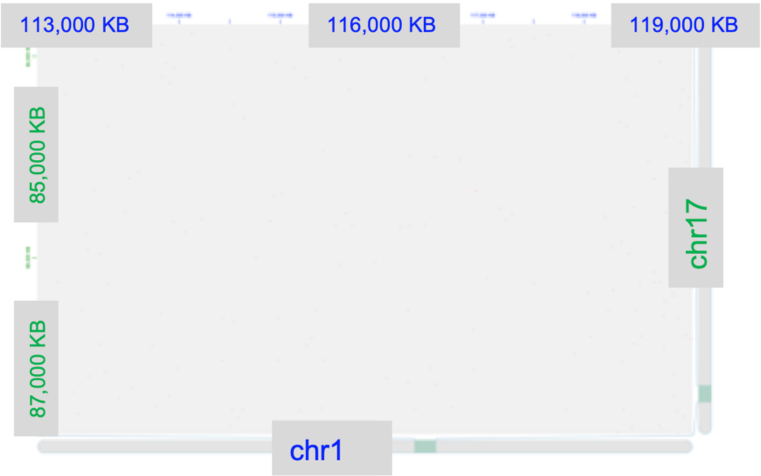


(D)

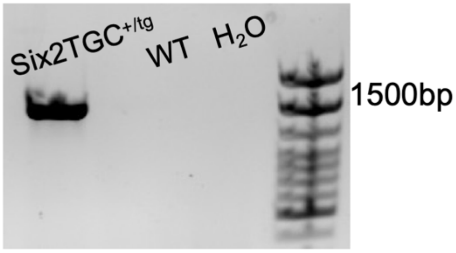

(E)

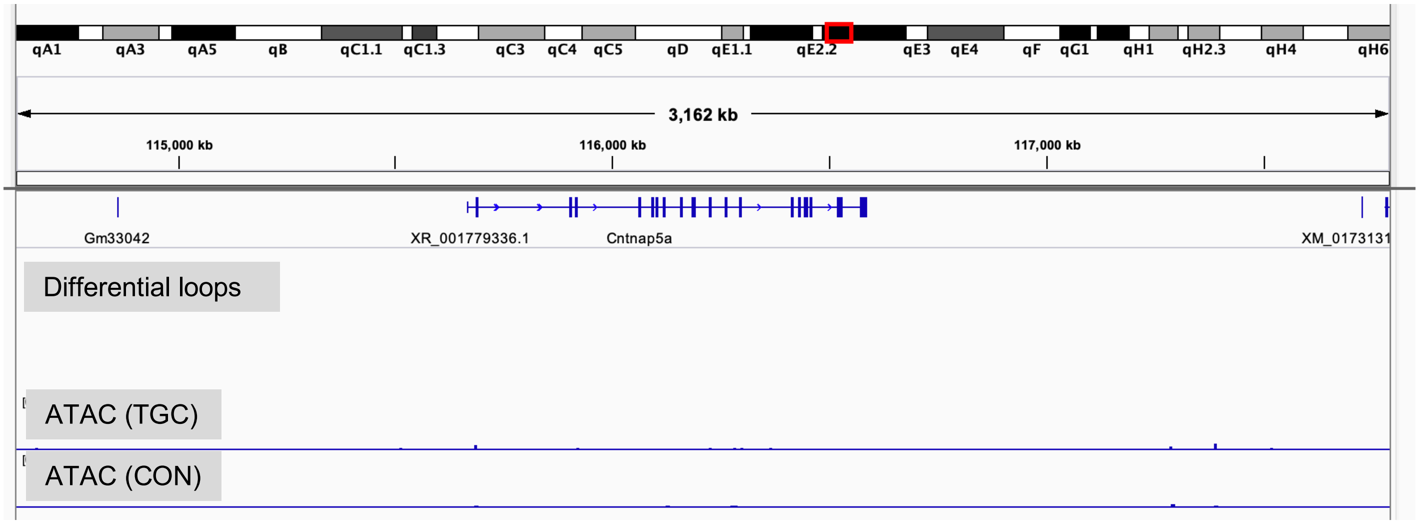


(F) 29.5MB view of chr1 centered on *Cntnap5a* (115MB)

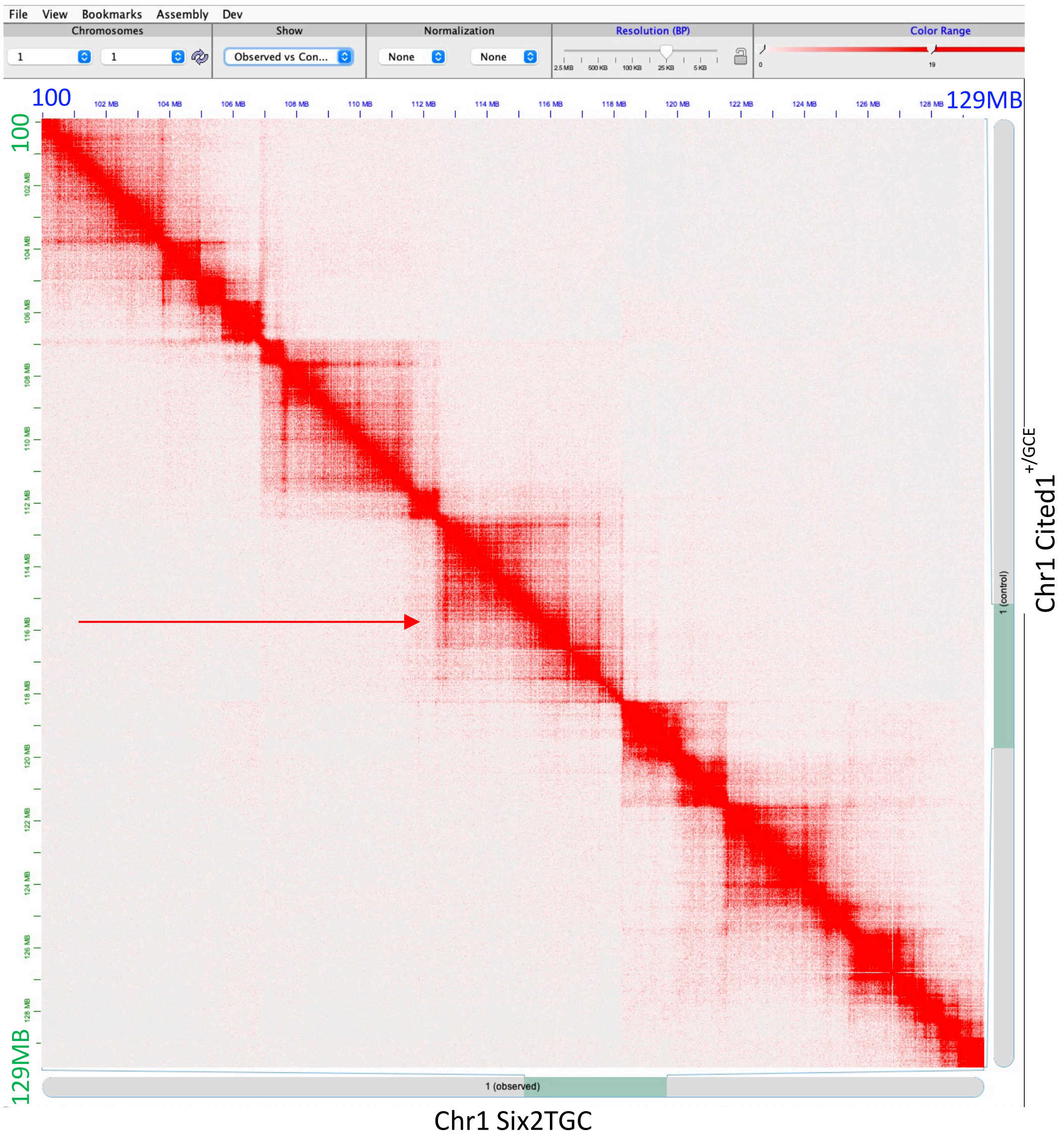

(G)

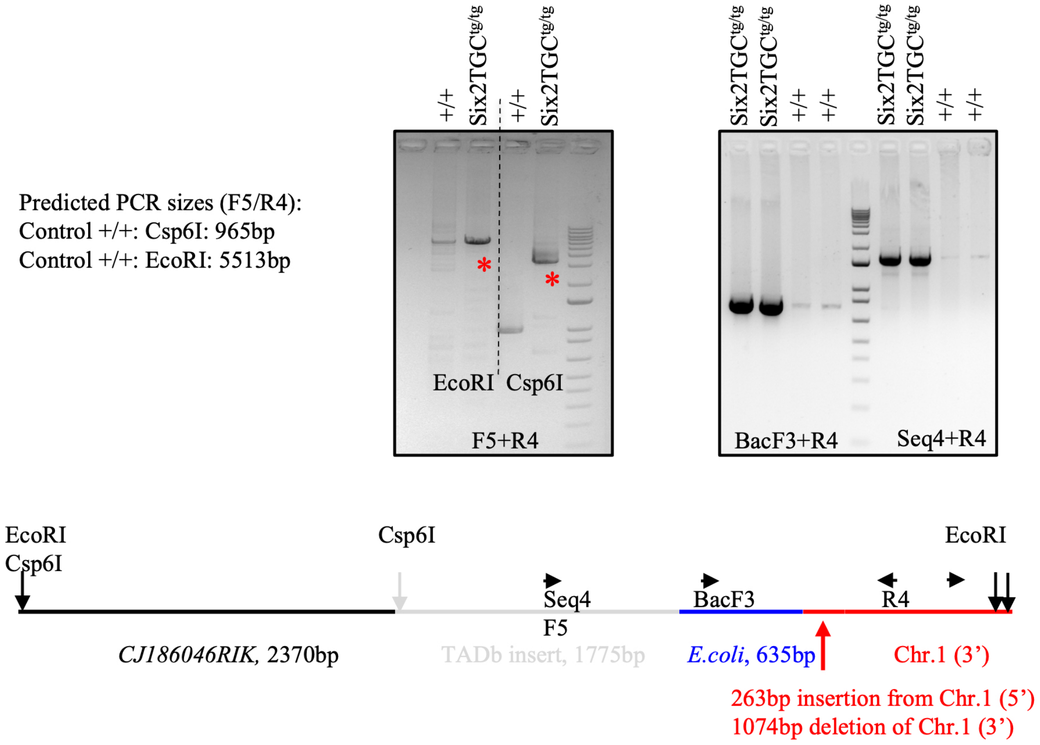


(H) Sequence of PCR product (F1+R1)

Red: GM41645

Black: CJ186046RIK

NNNNNNGNCNCNTGGCTGTCTTCCTCTNNATATTAATTCCTTCTCCTNATTATGGAATAACCCTGCTGTGCTGGAA
CTCACAGTACCTAGGCTTCTGCATCCAGTTATTCACCATAGAAAAGTCCTTAGCCAAAGGAAGAGAGAGATCCAAA
GTCTGACTTGTGGCTCTACCTATGGAAGGGAAGCTACAGTTTTTTCTTTGTGCAATTGAGAAGTATCAGCTCGTCT
GGGGTATCACATGGACTAGACTAGCTCTGGCTCTGGCTCTGGCTTCAGCCACTGGGACCAACAACCCATTGCTCTT
GCTGCCCTTCTAACTCATATCACCCTCTCACGCCCTTCACTCTTCTTGCCACCTTAGATTCGTGAGTTCTGTGTTT
ATAAATGTCTGTGCACATGGATTTTTTTCCTTCCCAGTCTCCTTCTCTCCTCGTTATACCAGTTGCCAAAAGTCTA
ATCCTTTTCAATATTTAATCTTCCACTACTTCATACCTCTACTCATACTAGCCAGTAGTGATTGAAACAAAGAAAA
ATAATTCACATTCGTGTCCCATAATTGTTTTTAATTCCATAGTATCAGCTAAGTTTTTTTTTTTAAAACAAAAGCT
TTTAAAAGGAAATTTACAATGATGATTGGTTTAAATGCGAAGCTGCATCAATATTGTGACGAGATCAATAATAGGA
TGTGAGTGGTTGCTGGCATATTGATCACATGTGTGAACAGTGAGTGATTGCTGGCATACTGATCANANATGTGAGC
NGNTTTGGACTGAGTACCANCTCCNAAGGAAANTGGGGTAAACGCNTACTCTGANTCTAGCCNGCCTCCANCCTCT
GGTCTTGAACCTANCATTTACACTACANCNTANCNTCCANCNNANNNCN

(I)

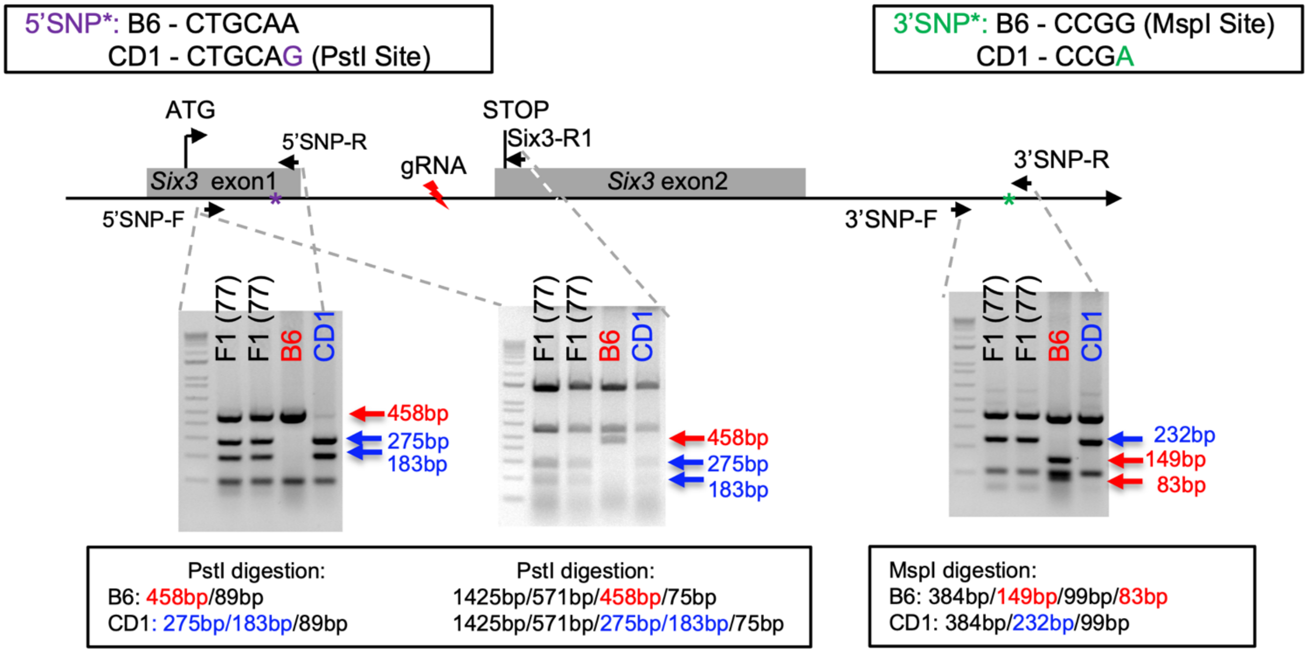


Genotyping of Six2TGC[77]-F1 mice with 2 SNPs (one in exon1, the other at 1975bp 3' of exon2) shows that the exon2 of transgenic *Six3* gene is deleted, while exon1 still present.

Primer sequences:

5'SNP-F: GAAGAGTTGTCCATGTTCCAGTTG
5'SNP-R: GAGTTTCCACCCGTTTCCATTT
Six3-R1: GAAGAGGAGCAGGGTTACGAAGAG
3'SNP-F: GCCAGATCTCTCTTTTACCTAGTG
3'SNP-R: CAGTCCCACTGTAGATCAGCATC

(J)

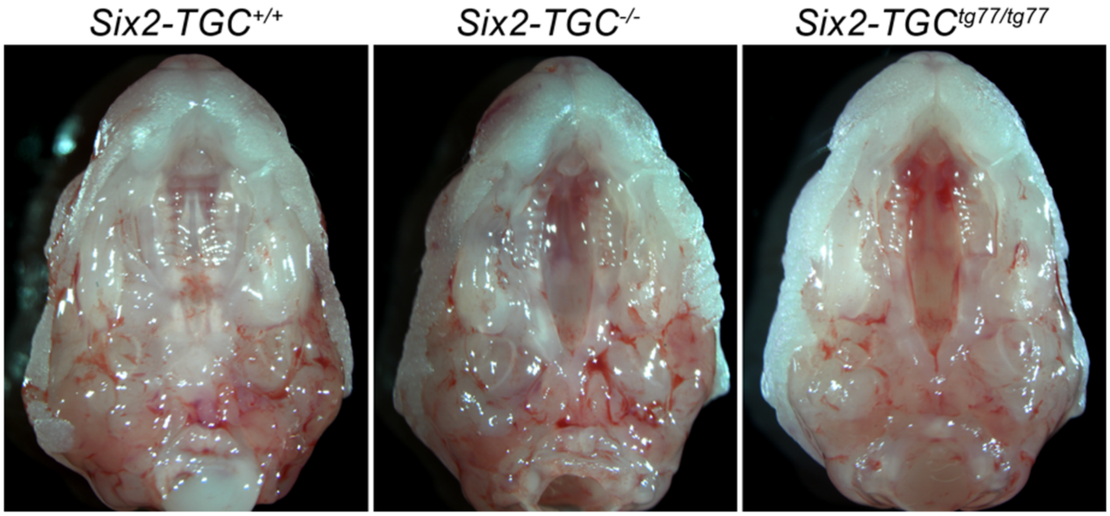

**Supplemental Figure S1. Supplementary analysis of *Six2TGC* allele architecture.** (A-B) Reports from TLA analysis performed by Cergentis (Utrecht, Netherlands;[(1)]) identified transgene integration within a 75kb span on chr1 located in a large intron of *Cntnap5a.* Whole genome overage plot (A) identifies signal for the endogenous chr17 locus and putative chr1 integration region, as well as (B) sequencing confirmation of the presence of the vector backbone fused to the BAC boundaries. (C) Juicebox visualization of chr17-chr1 interchromosomal contacts in control reveals paucity of interactions (compare with Figure 2A). (D) PCR screening of candidate BAC breakpoints amplified the 5' chr1-transgene junction (E) ATAC, Hi-C and RNA-seq data detect no differential transcripts, ATAC peaks, or novel/disrupted Hi-C loops. (F) 29.5 MB view of the chr1 from the *Cited1*$^{+/GCE}$ (green, red arrow points to the integration site) against the same region in Six2TGC (blue). (G) PCR strategy identifying architecture of 3' BAC integration site. (H) PCR product sequencing of BAC-BAC junction within *Six2TGC*. (I) Genotyping PCR strategy for identification of the exon2 deletion within the BAC for transgenic CRISPR founder line #77, versus the endogenous locus. (J) Cleft palate defect morphology for wild-type as well as homozygous animals from the original (Six2TGC) and CRISPR-modified line (denoted tg77).

Supplemental Figure S2

A

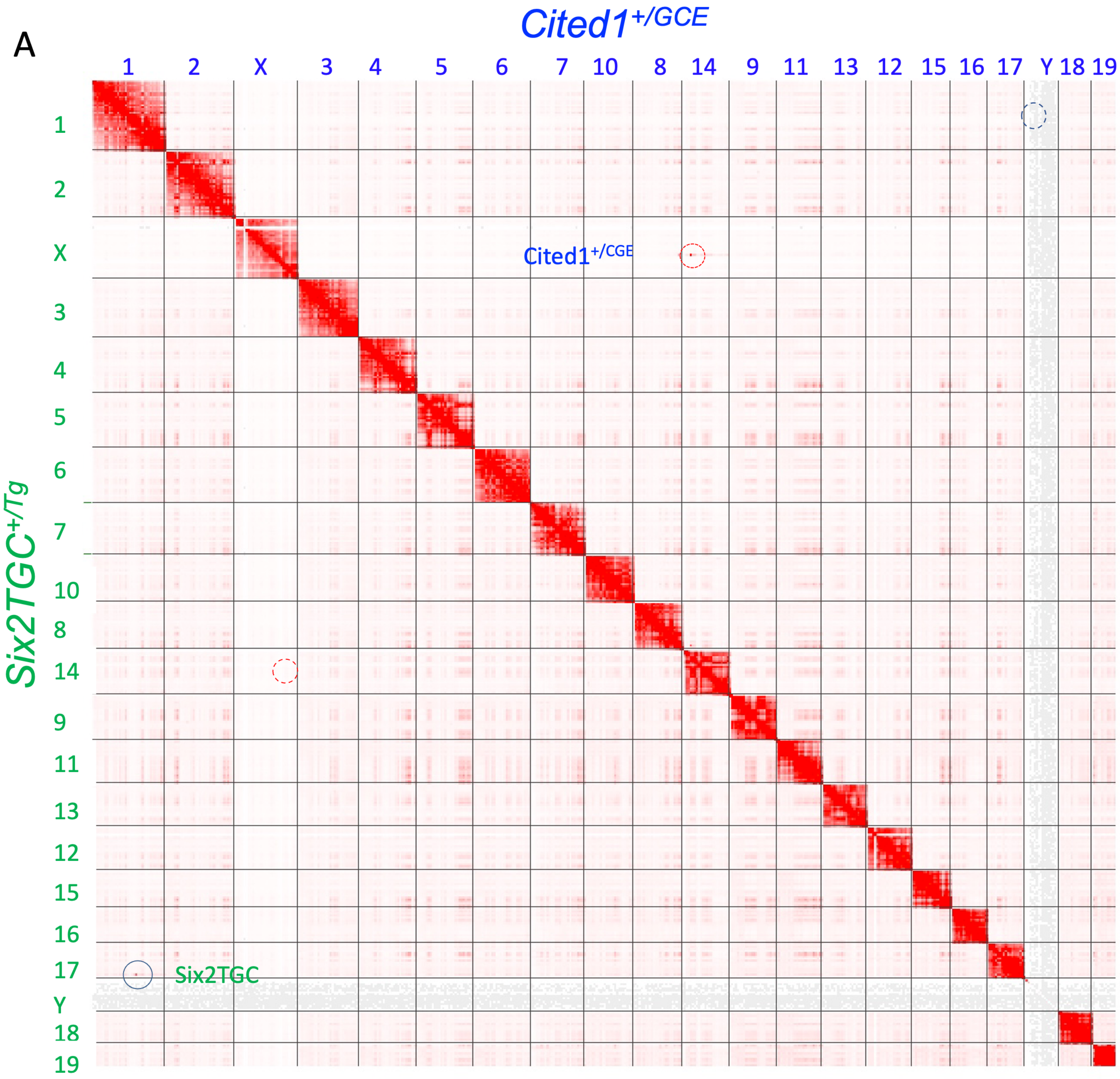

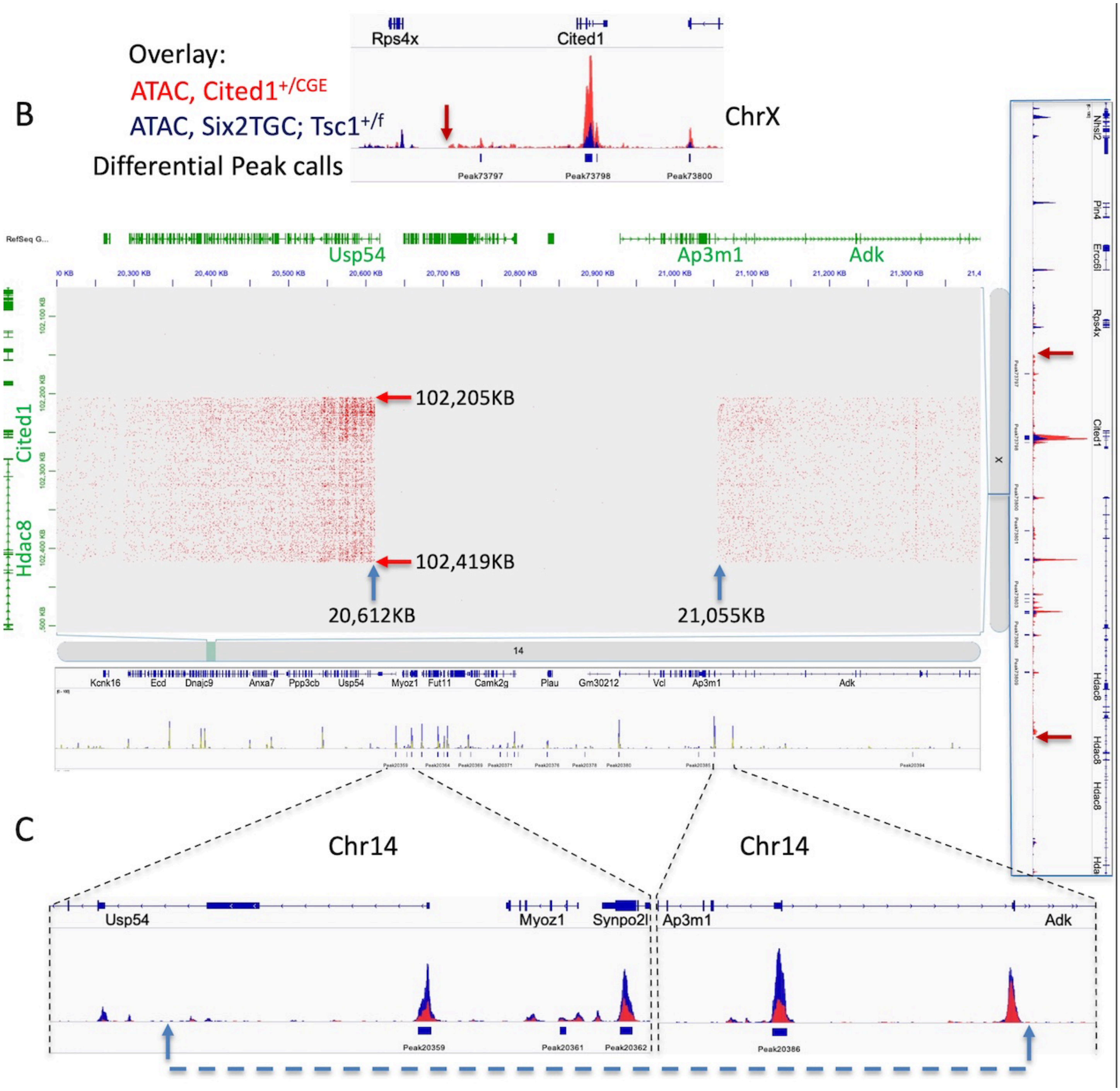


**Supplemental Figure S2. Cited1 transgene integration analysis.** (A) Alignment of all chromosomes from *Cited1*$^{+/GCE}$ (blue) against Six2TGC (green). Chr1/17 interactions in *Six2TGC* (blue dashed circle) are seen as an intense red dot which is not present in the reciprocal region of *Cited1*$^{+/GCE}$. Conversely, chrX/14 interactions are visible in *Cited1*$^{+/GCE}$ (red dashed circle) but not in *Six2TGC*. (B) Alignment of chr14 on the X-axis against chrX on the Y-axis, both from *Cited1*$^{+/GCE}$ , visualized in Juicebox, and IGV visualization of ATAC and differential peak calls in selected regions of both chromosomes. Differential ATAC peak calls with an overlay of the actual ATAC data shown on top for the *Cited1* region. The *Cited1*$^{+/GCE}$ tracks in red, the *Six2TGC; Tsc1*$^{+/f}$ in blue. The margin of the BAC shown by red vertical arrow. Note that four-fold more signal is coming from *Cited1*$^{+/GCE}$ chrX relative to *Six2TGC; Tsc1*$^{+/f}$*.* On the Y-axis, differential ATAC peak calls with an overlay of the actual ATAC data for the entire BAC. The width of the BAC is marked with red arrows which includes a portion of *HDAC8* and is present in four-fold excess as compared to *Six2TGC; Tsc1*$^{+/f}$. (C) Below the JuiceBox image we show ATAC-seq data and differential calls of chr14 region where the BAC integrated in *Cited1*$^{+/GCE}$. The large gap seen in

JuiceBox identifies a ~0.5MB deletion in chr14 (marked by vertical blue arrows), verified by differential peak calls in ATAC seq datasets spanning the gap and by a 1:2 ratio of peak height favoring the *Six2TGC* DNA.

**Supplemental Figure S3.**

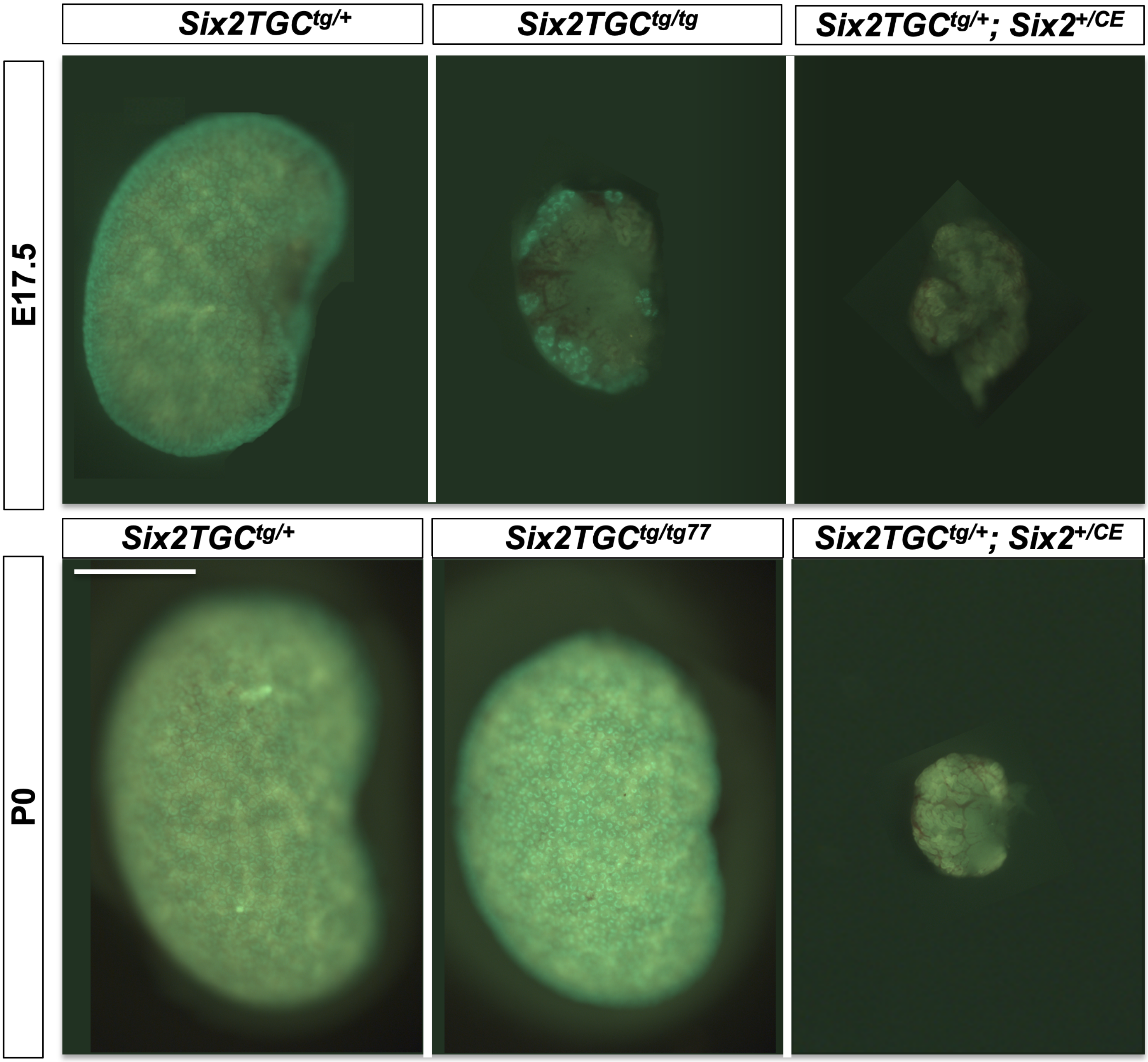


**Supplemental Figure S3. Kidneys from original and modified *Six2TGC* mice.** Surface GFP visualization at E17.5 and P0 for the indicated genotypes. All are taken at the same magnification as *Six2TGC*$^{+/tg}$. Scale bar denotes 1mm.
